# Time-varying nodal measures with temporal community structure: a cautionary note to avoid misinterpretation

**DOI:** 10.1101/659508

**Authors:** WH Thompson, G Kastrati, K Finc, J Wright, JM Shine, RA Poldrack

## Abstract

In network neuroscience, temporal network models have gained popularity. In these models, network properties have been related to cognition and behaviour. Here we demonstrate that calculating nodal properties that are dependent on temporal community structure (such as the participation coefficient) in time-varying contexts can potentially lead to misleading results. Specifically, with regards to the participation coefficient, increases in integration can be inferred when the opposite is occuring. Further, we present a temporal extension to the participation coefficient measure (temporal participation coefficient) that circumnavigates this problem by jointly considering all community partitions assigned to a node through time. The proposed method allows us to track a node’s integration through time while adjusting for the possible changes in the community structure of the overall network.

## Introduction

Quantifying the dynamics of a network often utilizes a multilayer network approach (Kivelä et al. 2014) of temporal ordered “snapshots” consisting of connectivity matrices through time (i.e. temporal network theory; Holme & Saramäki 2012). This approach answers questions about how nodes, edges, and communities in a network fluctuate over time. In recent years, such temporal network approaches have increased in neuroimaging (Shine & Poldrack, 2018). To generate knowledge about the underlying network, we also require that the temporal network measures can be mapped back to, or interpreted in terms of, the phenomenon they are modelling.

There are many metrics available for quantifying topological features of nodes within temporal networks. Some measures are temporal extensions of static measures (e.g. TempoRank is a temporal extension of PageRank (Rocha & Masuda 2014)). Others apply static measures to each time-point (e.g. Bola & Sabel 2015 found changes in rich club coefficients applied to multiple time-points). When applying static measures in a temporal network context, it is essential to ensure that the interpretability or clarity of the measure is not changed or distorted when used through time.

The participation coefficient (PC) is an example of a static network measure used in time-varying contexts by applying it to multiple time-points. Briefly, the PC quantifies the diversity of a node’s connections to other nodes across a community partition (Guimerà & Nunes Amaral 2005). The community partition groups different nodes based on a grouping property (e.g. modularity, when tightly connected nodes form modules (Newman & Girvan 2004)). Importantly, the PC for any given node is relative to the community partition used to calculate it (Figure 1A): if the community partition changes, then the participation coefficient may also change. In the two examples in Figure 1A, the shaded node has the same edges, but the communities are different, entailing that the participation coefficient changes.

**Figure 1:**
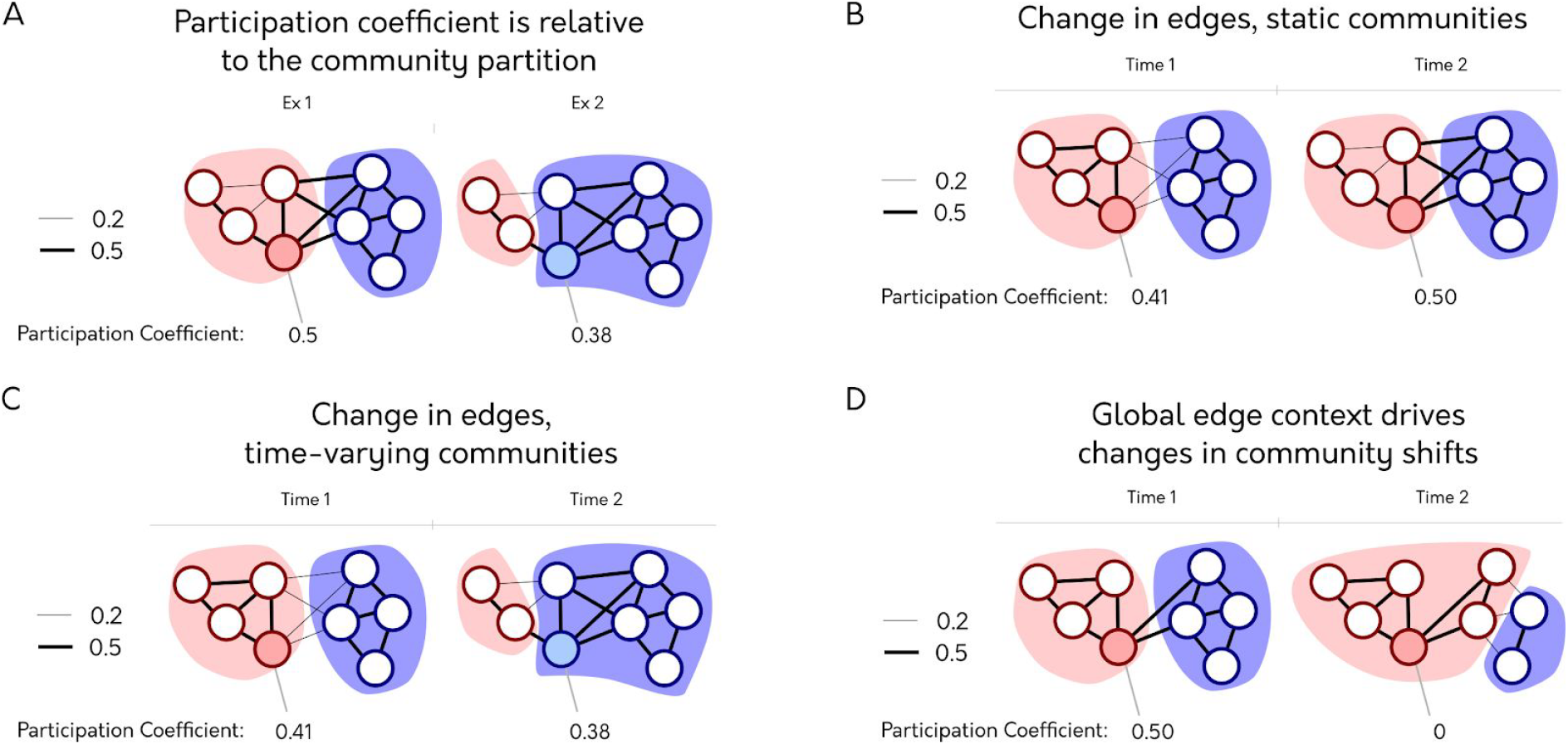
Different ways to calculate the participation coefficient through time. The participation coefficient (PC) for the shaded node is below each network/temporal snapshot. The border of each node and the coloured backgrounds show the assigned community of the node. In this example, there are two different edge weights possible. (A) Two examples of the participation coefficient, illustrating how the measure is calculated relative to the community partition. (B) An example of PC calculated when applying a static community template across multiple temporal snapshots (PC_S_). (C) An example of PC calculated when applying a temporal community partition to multiple temporal snapshots (PC_T_). PC values between time-points can not be directly compared due to differences in community structure obtained for each time-point. (D) An example showing that changes in community partitions over snapshots may result from changes in edges that do not directly connect to the node of interest. The difference in community partition is the result of changes in the nodes in the blue community. These changes affect the PC of the shaded node despite the fact that there was no change in its connectivity.

Community partitions can also be calculated through time. Temporal communities can merge, split, disappear and reappear through time (Granell et al. 2015). In the brain, the community structure has been shown to change in response to task and cognitive demands (Vatansever et al. 2015; Braun et al. 2015; Thompson et al. 2019).

Considering the wish to apply PC through time and that temporal communities in the brain fluctuate, it is understandable that many previous studies did both (e.g. Betzel et al. 2015, Shine et al. 2016, Pedersen et al. 2017, Tanimizu et al. 2017, Xie et al. 2018, Fukushima et al. 2018; Fukushima & Sporns 2018; Shine et al. 2018; Rizkallah et al. 2019). However, a problem with the interpretation of PC emerges when comparing two (or more) snapshots of a network with different community partitions. When community boundaries are allowed to fluctuate, as is the case with temporal communities, the participation coefficients calculated at different time-points may have different community-contexts.

Here we argue that calculating PC per time-point with a temporal community structure does not necessarily quantify its intended property of integration. As a result, the crucial link between the network measure and its concrete interpretation breaks down. The consequence of this that definitive conclusions cannot be drawn about what is happening in the brain. We demonstrate this problem on toy network examples and then using the resting-state fMRI data. Finally, we propose a new method - *the temporal participation coefficient* - that takes into account various community partitions calculated over time.

## Methods

### Data used and data accessibility

We used data from the Midnight Scan Club resting-state fMRI (Gordon et al. 2017) that is publicly available on openneuro.org (ds000224). The data is available after both preprocessing and denoising steps have been performed (see Gordon et al. 2017 for details of these steps). The data consists of ten subjects that underwent ten resting-state fMRI sessions. One subject was excluded due to substantial artefacts. We extracted time series from 200 functionally-defined parcels (Schaefer et al. 2018).

### Time-varying connectivity estimation

We estimated time-varying connectivity using weighted Pearson correlations with weights based on the Euclidean distance between time-points (Thompson et al. 2017). Briefly, the method creates a T×T matrix containing the Euclidean distance between all nodes at each time-points. At each time-point, the corresponding row of the distance matrix becomes a weight vector for the connectivity estimates. The weights (W) are first converted into a similarity matrix (1-W) and then scaled between 0 and 1. This vector is then used as the weights in the covariance matrix to create a weighted Pearson correlation. While the sliding window approach uses “temporally nearby” time-points to support its connectivity estimates when estimating the covariance, this method uses “spatially nearby” time-points to support its connectivity estimate (see Thompson & Fransson 2018 for illustrations). We selected this method of functional connectivity estimation because it performs well (2nd place) at tracking a fluctuating covariance through time in simulations (Thompson et al. 2018). The method that was ranked first - jackknife correlation - was not chosen here because it calculates the “difference in connectivity” which has a different interpretation than the usual methods.

Prior to calculating the Louvain community detection and the participation coefficient, all edges below 0 were set to 0.

### Quantifying community structure

We calculated the temporal communities using the Louvain algorithm (Blondel et al. 2008) with the resolution parameter of 1. Temporal consensus clustering was performed by assigning communities at time t-1 with the same label as the community at time t that had the smallest Jaccard distance (Lancichinetti et al. 2012). We also calculate static functional connectivity using Pearson correlations and a static community partition with the same parameters as the temporal communities.

### Static participation coefficient (static PC)

The participation coefficient is defined by Guimerà et al. 2015 as:

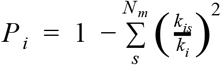

Where *i* is a node index; *N*_*M*_ is the number of communities. *k*_*is*_ is the within-community degree, and *k*_*i*_is the overall degree of node *i*.

### The participation coefficient through time with static communities (PC_s_)

When calculating the participation coefficient through time, with static communities, the equation is:

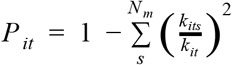

Here we see temporal subscripts for the participation and the degree of the nodes. Note, there is only one community partition used for all time-points.

### The participation coefficient through time with temporal communities (PC_T_)

The participation coefficient with temporal communities is:

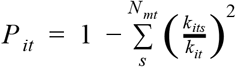

Above, we are summing over the communities N_mt_ which is the number of communities found at time-point *t*. This method uses a community partition calculated separately per time point.

### Temporal participation coefficient (TPC)

The temporal participation coefficient that we introduce is:

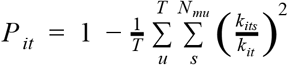

Here we have added a term, where all temporal community partitions are considered for each time-point. See the results section for the motivation behind the TPC.

We abbreviate the different participation coefficient methods as follows: “the static participation coefficient” (static *PC*), “the participation coefficient per time-point with static communities” (*PC*_*S*_), “the participation coefficient per time-point with temporal communities” (*PC*_*T*_), “the temporal participation coefficient” (*TPC*).

### Additional network measures

We also related the different participation measures to different network estimates throughout the article to help interpret what the changes in participation means for the network. The additional measures we used were: (i) flexibility (the percentage of times where a node changes its community, Bassett et al. 2011); and (ii) within-module degree z-score (*z*; z-score of node’s degree within its community, Guimerà & Nunes Amaral 2005). We calculated *z* in two ways: using static communities (*z*_S_) and using temporal communities (*z*_T_). We combined the different *z* estimates with their methodological participation counterpart (i.e. *z*_*S*_ is used with *P*_*s*_).

### Illustrating the methodological choices do not impact the conclusion

To ensure that our concern about PC_T_ generalizes across different methods, we replicated part of the analysis (Figure 5A) by changing some methodological choices: (i) the time-varying connectivity estimation method and (ii) the community detection algorithm. As an alternative time-varying connectivity estimation method, we used the multiplication of the temporal derivative (MTD) method (Shine et al. 2015). This method multiplies the temporal derivatives of each pair of time series and applies a smoothing parameter that averages over +/- 4 time-points. When changing the community detection algorithm, we applied TCTC (Thompson et al. 2019, with parameters: ϵ: 0.5, σ: 5, τ: 5, κ: 1). Since this community detection is multilabel, we created single labels partitions by applying the Louvain algorithm (resolution parameter: 1) to drive each node into a single community.

## Results

### Different reasons underlying changes in participation coefficient through time

Here, we illustrate the potential problem introduced when calculating the participation coefficient through time on some toy network examples. Consider a time series of participation coefficients when the community partition is static (Figure 1B). For the two different temporal snapshots, there is a change in the edges of the shaded node, which changes the participation coefficient of that node. Specifically, in the second snapshot, the connections of this node have become evenly distributed across nodes in all communities. We can easily relate the two *PC* values for the two different snapshots to each other, and it makes sense to interpret the increase participation coefficient as an increase in the node’s interaction with communities outside of its own.

If instead, the community partition varies over time (Figure 1C), the changes in edges lead to the shaded node being classed as part of the blue community instead of the red community. The node’s participation coefficient, in light of this change in community membership, is reduced. In the second snapshot, the participation coefficient has changed - it has decreased - not because the node decreased its participation (i.e. its role in the network). Instead, the node changed community membership when it increased its connection strengths with the blue community relative to the first time-point. Hence, the interpretation of a temporal series of participation coefficients as reflective of a change in intra-community connections is impeded by the extent the community structure is changing over time. That is, none of the *PC* values in a time-series can be directly compared to each other, nor can we directly translate the abstract measure to the external phenomenon.

A possible objection to this criticism of PC_T_ is that the temporal communities are calculated on the edges themselves, entailing an interconnection between the community-context and edge-context of a node. This objection does not adequately take into account how communities are calculated. Communities take into account the “global edge context” (i.e. all edges in a network and how they relate to each other) whereas the participation coefficient only considers the “local edge context” (i.e. all edges connected to one node). There is no necessary relationship between these two (exemplified in Figure 1D). A node’s strength can increase with no effect on the community partition. Alternatively, a node can change its community assignment with no change to its own edges.

### Misleading network-level interpretations of PC with temporal communities

We have demonstrated that community-context affects the nodal participation coefficient when quantified at multiple snapshots. The measures are applied correctly according to their mathematical definitions, which raises the question about why this is of any importance. Here we show how PC_T_ can cause misinterpretations regarding the property that participation is generally used to quantify.

What property is the participation coefficient trying to identify? In its introduction by Guimerà & Nunes Amaral 2005, they claim that “[t]he participation coefficient *P*_*i*_ measures how ‘well-distributed’ the links of node *i* are among different modules” (p.g. 897). This property of “well-distributed” edges has been interpreted as the *integration* within network neuroscience when applied to static functional connectivity. For example, Bertolero et al. (2015) interpreted their results regarding PC as: “nodes with high participation coefficients integrate information and coordinate connectivity between communities” (p.g. 2). Likewise, Power et al. (2012) stated that the participation coefficient interpretation relates to spanning multiple different systems: “If a node has a high participation index […] we infer that such nodes likely have access to a variety of types of different information processing represented among different systems” (p.g. 808). PC_T_ kept the same interpretation when applied in time-varying contexts. Shine et al. (2016) stated that they identified “functional states that maximize either segregation into tight-knit communities or integration across otherwise disparate neural regions” (p.g. 544) where the integration was calculated using PC_T_. Thus, from its network science origins to functional connectivity in network neuroscience, to time-varying connectivity, the participation coefficient was consistently considered to quantify the integration of nodes in a network.

We will now show that PC_T_ does not always quantify integration, as described in the literature. Consider the toy network demonstrated in Figure 2A. Here we have two different time-points where the community-context changes if considering temporal communities. We have selected nodes from this network and identified when PC_S_ and PC_T_ would exhibit increased integration, segregation or no change in regards to the previous time-point (Figure 2B). Note how PC_T_ assigns high participation and thus interpreted as having higher integration, to the node marked in blue in Figure 2B. The difference between the network at Time 1 and Time 2 is that the blue community has split into two smaller communities. Following the interpretation that integration represents the information processing across different communities (e.g. quote by Power et al. 2012 above), it is hard to consider a split in community structure as increased integration — in this example, PC_T_ is failing to quantify integration. Likewise, the temporal community of the red node has extended in the second time-point (Figure 2B, Time 2). This extension of the community means that the red node is sharing similar information with more nodes than usual, yet PC_T_ will ascribe a low score entailing that there is less integration at the second time point. In both these examples, PC_S_ provides the opposite values. Thus, high PC_T_ can lead to the incorrect interpretation that more integration is occurring in the network. Note, there is nothing mathematically wrong with each PC_T_ estimate, the problem lies when contrasting PC_T_ estimates with each other and inferring changes in integration from such a contrast.

**Figure 2.**
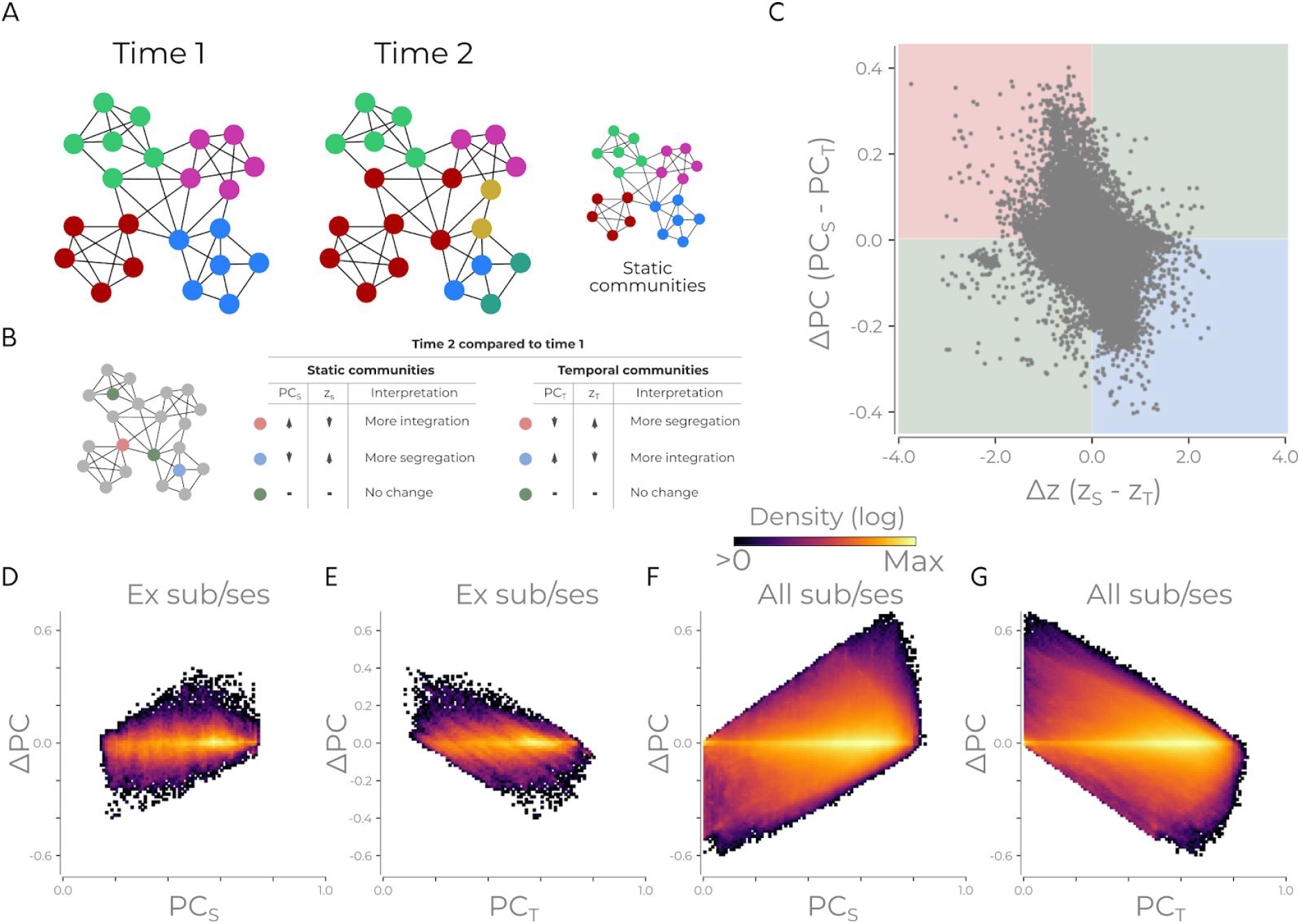
The different PC calculation methods lead to different interpretations of the network’s organization. (A) An example network consisting of multiple communities for two time-points. The figure shows both a temporal community partition and a static community partition. (B) Different types of nodes from A are highlighted. The accompanying table states how these nodes at time-point two will be quantified relative to time-point 1 with static or temporal communities for the participation coefficient (PC) and within-module degree z-score (*z*). (C) The difference in PC and *z* for nodes in the static and temporal for all nodes/time-points for one subject/run in the MSC dataset. The red, blue and green quadrant corresponds to the interpretations found in B. (D) Density plot showing the difference in PC versus PC_S_ for the example session/subject in C. (E) Same as D but the difference in PC versus PC_T_. (F) Same as D but for all sessions/subject. (G) Same as E but for all sessions/subjects. All density plots have a logarithmic colour scale.

Do situations where PC_S_ and PC_T_ have flipped interpretations occur on empirical data (i.e. situations like Figure 2B)? To test this, we calculated the difference between PC_T_ and PC_S_ and between z_T_ and z_S_ on an example subject’s resting-state fMRI data (Figure 2C). In Figure 2C, the coloured quadrants represent situations similar to Figure 2B’s node examples. Here the red quadrant indicates that they would have “more integration with static communities, more segregation with temporal communities” and vice versa for the blue quadrant. A large portion of the nodes (example session/subject: 75.59%; all subjects: 66.08%) end up in the red and blue quadrants, entailing that they have alternative interpretations about what is occurring in the network when using PC_T_ and PC_S_ methods.

For completeness, Figure 2D-G shows when the absolute difference of PC is high, then one of the methods with have high PC and the other low PC. The difference between PC methods is related to PC values of both methods for both an example session/subject and all subjects in the dataset. This relationship is expected as PC values are between 0 and 1. However, the differences between methods scale with PC magnitudes. This shows that the divergence between the methods is occurring at time points that are offering opposite interpretations about the node’s role in the network.

When contrasting two time-points in a multi-layer network with different communities per time-point, it is impossible to determine, from the values themselves, when the node is “more integrated”. Thus, given the participation coefficient’s long history with being used to quantify the amount of integration in the brain, using PC_T_ this way will lead to misleading results. Importantly, PC_S_ does not suffer from this problem as it uses the same reference community. However, the temporal community information is discarded, leading us to our possible modification of PC to allow for temporal communities.

### PC in relation to all temporal communities

Given the substantial evidence for temporal changes in community structure, there is an understandable desire to calculate the participation coefficient with fluctuating community structure. We present a possible solution to the problem outlined above: the temporal participation coefficient (TPC). The crux of the problem is that the participation coefficient of a node is relative to the community partition. If instead, each participation coefficient estimate considers all possible community partitions that the node has been assigned, then the participation coefficients will be comparable across time-points as each estimate is now relative to the same community context (Figure 3). This solution calculates the local edge context at a time-point with all possible community contexts, weighted by the frequency of each community. Then it considers how a node is participating relative to the possible community structure it can have. In Figure 3, each TPC estimate is calculated relative to both community contexts, entailing that the shaded node at the second time-point has more participation compared to the first time-point. As both time-points have their different edge-contexts calculated relative to the same set of community partitions, these values can now be compared.

**Figure 3:**
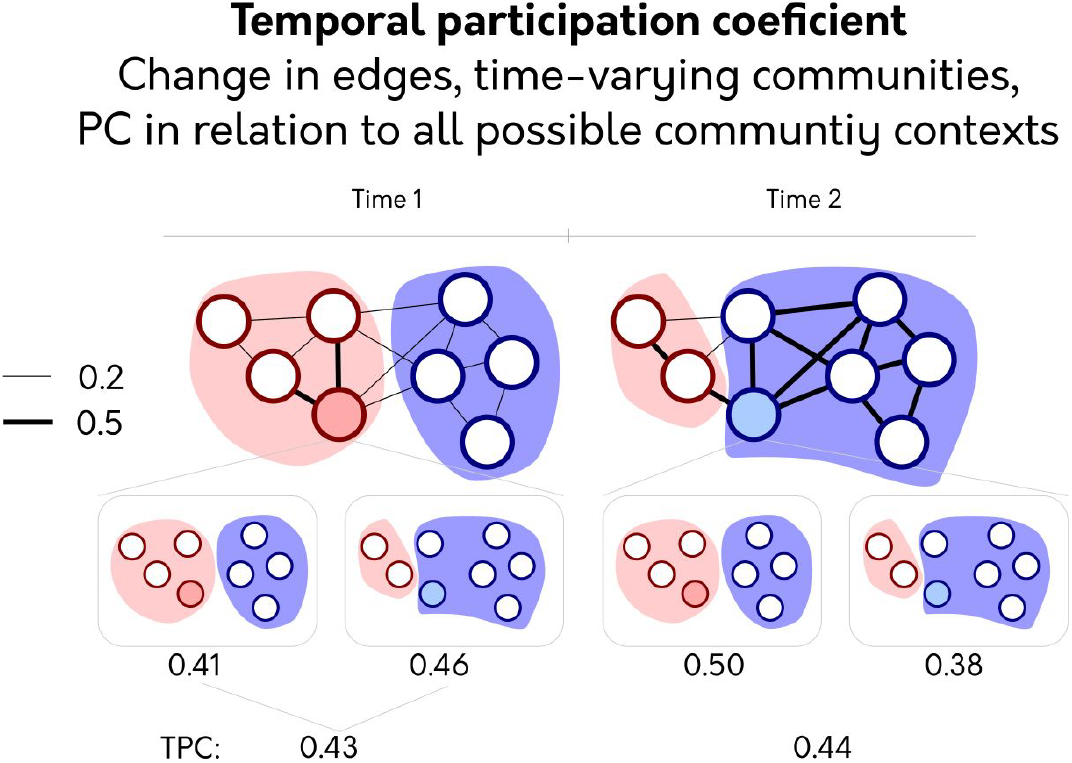
The temporal participation coefficient (TPC). To calculate the TPC, PC is calculated per time point, taking into account all possible community contexts the node can be in. PC value calculated for each possible temporal community context is shown under each network. These are then averaged together. This fix makes PC of the shaded node comparable across time-points as they have been compared against the same community contexts.

The motivation behind TPC is to have all time-points compared to the same community context (like PC_S_) but to retain the temporal information inherent in community fluctuations (used in PC_T_). Contrasting time-points’ participation from different community contexts, as shown above, can lead to misleading interpretations. To rectify this and still utilize the time-varying community structure, we calculate the degree at each time-point with regards to all possible community partitions. Perhaps this logic seems counter-intuitive as it entails community partitions that do not fully reflect the snapshot. However, this is also the same logic as PC_S_, as it applies a static community partition to each time point that does not fully reflect the community structure at each snapshot. The only difference here is that TPC utilizes multiple community partitions instead of a single partition.

There is one crucial assumption when applying this method, namely that the community structure can recur again through time. This assumption means that the same or similar community partitions will be found at later time-points (i.e. a similar snapshot can occur at a later time). In a network like the brain, this is reasonable, and it is an assumption that is made in all psychological experiments where the task is repeated (see discussion for when this assumption breaks down). Without this assumption, then it becomes unfair to user earlier or later community partitions to analyse a node’s PC in a snapshot as if it was in another community.

TPC quantifies a node’s activity in relation to all communities it could potentially be in. Consequently, if a single large edge at t_1_ merges two communities, and, at t_2_, only that edge changes and the community splits, TPC will assign higher participation to t_1_. Since all time-points use the same community information, it is mathematically impossible for any time-point that decreases all of its edge weights to get higher participation — this guarantee is not possible for PC_T_. Thus, TPC can utilize all community contexts but avoids the problematic applications shown for PC_T_ in the previous section.

### Nodes with high static PC change communities the most

We have presented a theoretical problem, illustrated how it could lead to differences in interpretation and presented a potential fix. We have yet to show that the extent of the problem affects data itself. It *could* be the case that there is no difference when applying TPC versus PC_T_.

We begin by asking the question: how are nodes with high static PC affected by the temporal community partitions? If nodes with high participation have little change in their community context, the problem we raise may be redundant. To clarify this, we compared the flexibility with the static participation coefficient (Figure 4AB). Here we see that nodes with high participation also increase their flexibility. If nodes with high participation always remained in the same communities, calculating PC with temporal communities would be less problematic. Nodes that switch temporal communities have high static PC; thus, it seems concerning for participation calculated through time relative to different community partitions. We have not proven which participation method should be preferred, but it shows that the community contexts are affecting nodes with high participation and will help us understand any differences between participation methods.

**Figure 4.**
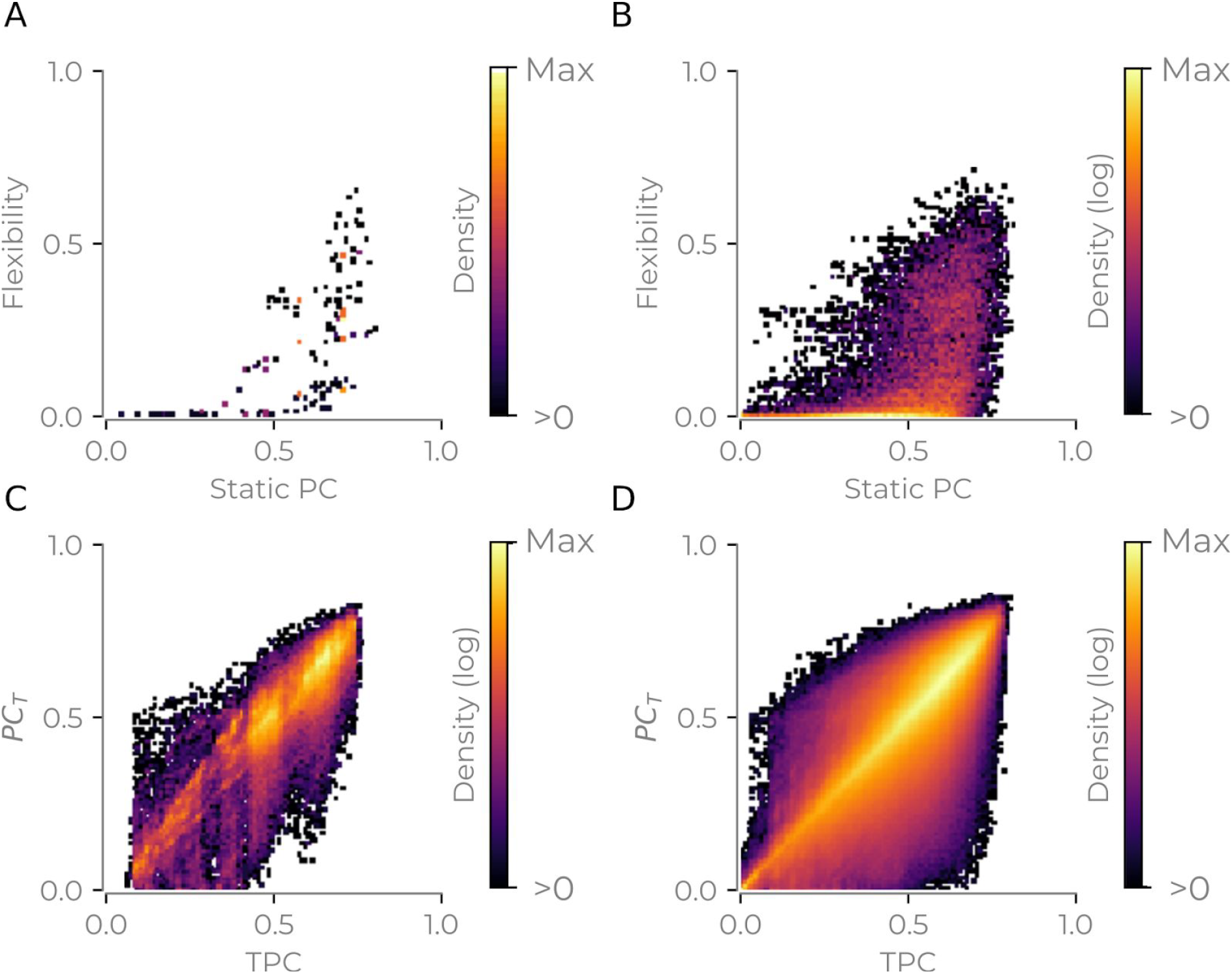
Differences between TPC and PC_T_. (A) Static participation coefficient versus the flexibility for one subject/session. (B) Same as A but for all subjects. (C) Temporal participation coefficient versus the participation coefficient per time point with temporal communities for one subject/session. (D) Same as C, but for all subjects. All colour bars show density on a 100×100 grid. Panels B-D show logarithmic range between colours.

### Divergence of the different methods

Now we contrast PC_T_ with the TPC to see whether they compute similar values or whether they diverge. First, we begin by considering all time-points and all nodes together. A heteroskedastic relationship between the two coefficient emerges (Figure 4CD, Bartlett test for heteroskedasticity: all subjects: *T* = 153221.9, *p* < 0.001; example subject: *T* = 1902.3, *p* < 0.001). This heteroskedastic relationship entails that, while both methods may identify points that have the highest participation, the relationship quickly breaks down. For completeness, Supplementary Figure 1 shows the relationship between both PC_T_ and TPC with PC_S_ similar to Figure 4CD. It can be observed that the extent of the heteroskedastic spread in Figure 4CD is largest for PC_S_ and PC_T_.

To quantify the extent to which the methods diverge, we considered two different questions: (1) do the time-series of participation coefficients correlate with each other?; and (2) If selecting the top *x*% of time-points to be marked as candidate temporal hubs, do the selections intersect? The time series of PC_S_ and PC_T_ did not correlate highly (Spearman rank (ρ): median: 0.30, SD: 0.21, min: −0.44, max: 1.0, Figure 5A), especially compared to PC_S_ and TPC (Spearman rank: median: 0.96, SD: 0.19, min: −0.80, max: 1.0, Figure 5B). TPC also did not correlate highly with PC_T_ (median: 0.32, SD: 0.21, min: −0.37, max: 1.0, Figure 5C). In sum, these correlations of the time series show that TPC and PC_S_ correspond the most with each other. However, while there is generally a positive correlation between TPC and PC_S_ time series, some of the time series differ quite radically (i.e. min correlation value is −0.80). This illustrates that these two methods can produce very different results for some nodes.

**Figure 5:**
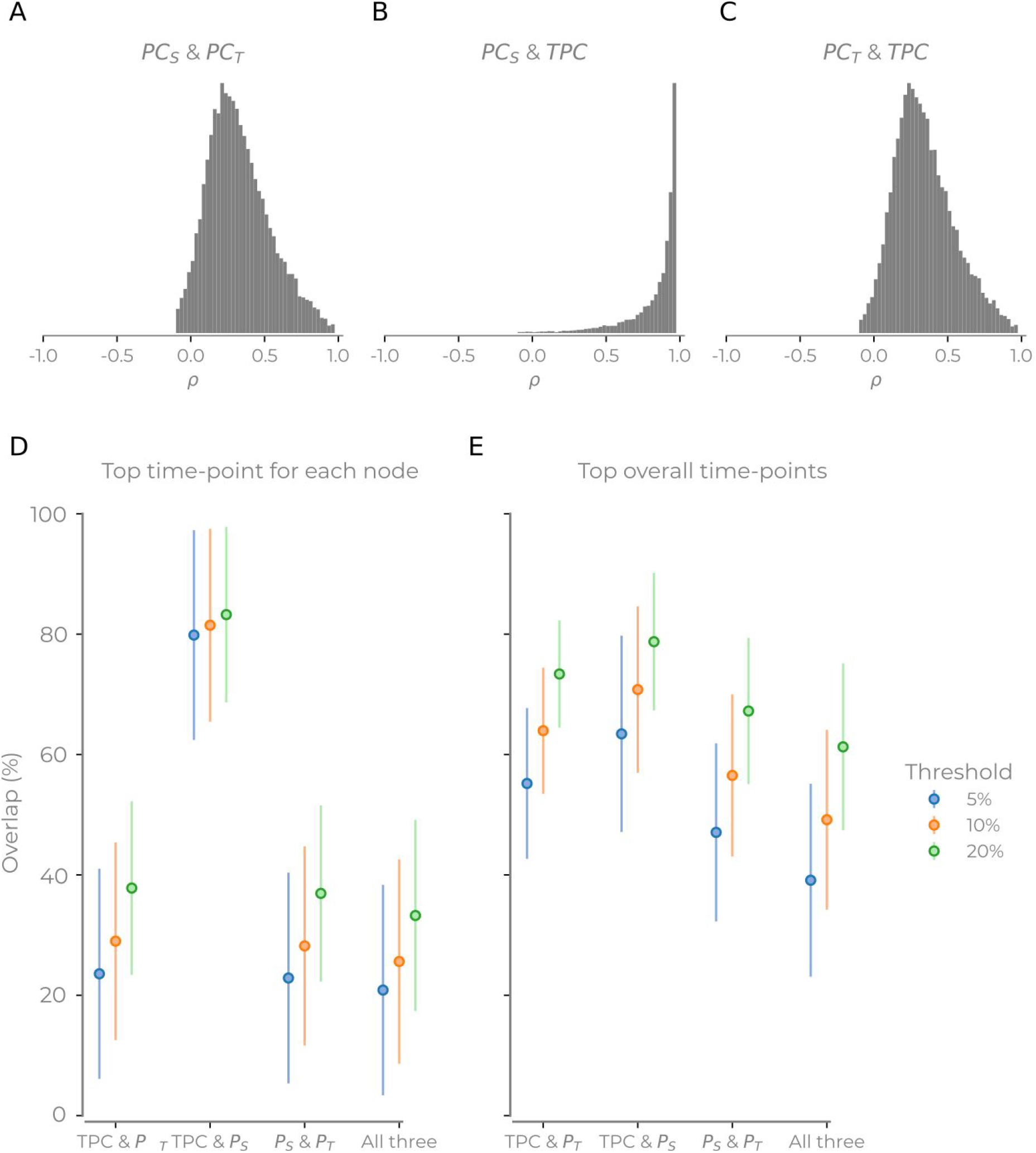
Comparison of the hub overlap for three different PC methods. (A-C) The histograms of correlation values for each time series for different participation methods. Histograms show all nodes, sessions and subjects. (D-E) The intersection of high participation coefficient from different methods. Here we see the intersection of each combination of three methods. (D) For each subject, the top x% time-point for each node. (E) For each subject, the top x% across all nodes and time-points. Error bars show standard deviation.

Taking the mean PC over time for the various methods, the correlation between the methods is high for all method combinations (PC_S_ & PC_T_: =0.91; PC_S_ and TPC: 0.91; PC_T_ & TPC: 1.0). This shows that the average PC over time is the same across methods. Importantly, the divergence of the methods is only when fluctuations are considered.

The correlation between PC time series calculated with different methods does not mean that the same nodes or time-points will get selected as candidate hubs as the correlation is only quantifying their covariance. For each method, we then identified the highest 5, 10 and 20 percent of values for both for the top time-points for each node (Figure 5D) and the top overall values (Figure 5E). If we try and find when each node has its highest participation, we find that PC_T_ has more unique nodes. When pooling all nodes and time-points together, the overlap of all three methods reached over 60% with larger thresholds but was under 40% for lower thresholds. This result shows that the choice of PC method matters. Finally, we also observed that the TPC and PC_S_ overlapped the most (reaching 80% of nodes in some instances and always over 60% when combining the paired and triple intersections). This overlap is reassuring for TPC as we know PC_S_ is a valid method. Moreover, the divergence that happens between the TPC and PC_S_ with static communities is due to the TPC utilizing the temporal community information.

Finally, we replicated Figure 5A by changing the time-varying connectivity method (to MTD) (Supplementary Figure 2A) and by changing the community detection algorithm (to TCTC) (Supplementary Figure 2B). The general pattern remains when contrasting Figure 5 with these other methods. The ordering of the peaks of all distributions follows a similar pattern. However the distributions changed in shape for PC_S_ and TPC, but the peaks of the distributions remained in the same order as Figure 5A, entailing that these methodological choices may diverge more. More importantly, there is no sizeable improvement with PC_T_’s correlations with the other methods, entailing that the interpretation problem with PC_T_ will persist to give misleading results. In sum, methodological variability does not induce the problems we have raised in this article.

## Discussion

We have outlined why PC_T_ can lead to misleading interpretations when contrasting different temporal snapshots. Further, we have proposed the temporal participation coefficient, which allows for across time-points comparisons without impeding the interpretation. Finally, we have also shown that these methods diverge in how much nodal time series correlate and which nodes will be considered hubs. The extent of the divergence between PC_T_, PC_S_ and TPC will depend on how much the communities fluctuate. The extent of the divergence will depend on both the parameters, time-varying connectivity method, community detection algorithm, and the ground truth but the methods do diverge. However, when changing both the time-varying connectivity method and community detection algorithms, were consistently PC_S_ and TPC the closest to each other, illustrating that PC_T_ diverges the most.

We are not challenging PC_T_ in all use cases; it is mathematically sound when applied to each time point. The problem arises when contrasting values from different time-points that are derived on different community vectors. If two snapshots are contrasted with PC_T_ and presented with their respective community differences each estimate can be useful for quantitatively understanding what is occurring in each snapshot and allows for the snapshots to be compare the time-points qualitatively (e.g. Figure 1D) which can facilitate understanding about the network — however, PC_T_ snapshots cannot be contrasted unless you alter the meaning of integration.

Measures of network neuroscience aim to increase our understanding about the organization of the brain. Our results show that PC_T_ can lead to misleading interpretations, especially when inferring a property such as integration. At the regional level, the time series between PC_S_ and PC_T_ correlated around 0.3 on average during resting-state (that is only sharing 9% of the variance). Our discussion has shown that it is unclear whether changes in PC_T_ occur due to communities splitting or via increased integration, and that is not possible to know what PC_T_ is quantifying based on the value alone. Hence, interpretations of PC_T_ definitively being a measure of integration when comparing multiple time points is, in our opinion, ill-advised. However, averaging over time points is possible for PC_T_, which multiple studies have done (e.g. in Shine et al 2016). This strategy is possible because it is no longer comparing time-points with different communities is possible and unproblematic (but loses the temporal resolution of the PC). In sum, we feel that for any quantification of fluctuations of participation through time should use PC_S_ or TPC, if changes in community structure are important for the research question at hand.

The solution we present, TPC, does not fit all possible use-cases. One limitation is that it can only be applied when the network can return to previous states (the recurrence assumption). Some temporal communities may only be possible after certain events have transpired - e.g. during a contagious outbreak, patients could form communities in the hospital. Using our proposed TPC fix on such a data set would entail that post-infection communities would influence pre-infection participation, which would be unrealistic. Furthermore, care would also be needed for any of the participation methods if a network bifurcates its community structure between entirely different states that have little or no topographic overlap as combining the network contexts may be hard to interpret. Thus, the proposed solution only covers networks which can theoretically return to similar states again, and restraint on the possible temporal community structure exists (e.g. anatomy). This assumption appears reasonable for networks such as the brain. However, quantifying variations in how nodes relate to their community assignments (e.g. Bassett et al. 2011) or using time-varying measures with static communities (e.g. PC_S_) may be more prudent analysis alternatives. The ultimate lesson here is that network measures need to be chosen based on knowledge about the system under investigation and the new information that the measures hope to gain.

Here the focus has been on temporal communities and its recent application within network neuroscience. However, this can also be a more general warning for such nodal measures that are relative to the community structure when applied in multilayer cases and may need to the fix we propose here).

We hope that this article highlights the problematic nature of quantifying temporal nodal measures relative to a fluctuating temporal community partition. We have offered one possible solution for this problem that utilizes temporal community information that does not suffer from similar issues regarding interpretation.

## Acknowledgements

WHT acknowledges support from the Knut och Alice Wallenbergs Stiftelse (SE) (grant no. 2016.0473, http://kaw.wallenberg.org). KF was supported by the Foundation for Polish Science, Poland (START 23.2018). There are no conflicts of interest to declare.

---

**Supplementary Figure 1:**
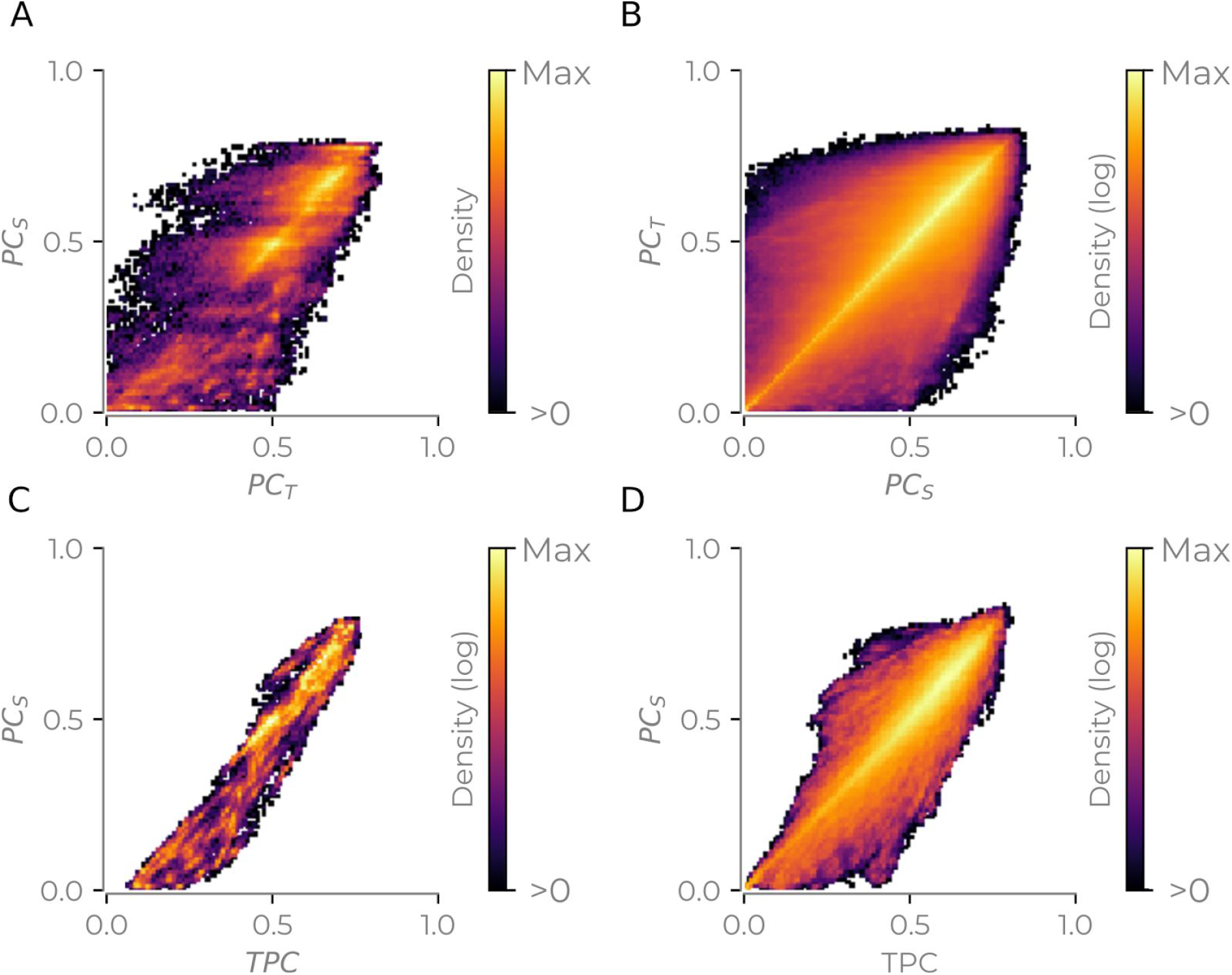
Same as Figure 4CD in the main text but showing the relationship with PC_T_ and TPC with PC_S_. A and C show example subjects. B and D show for all subjects.

**Supplementary Figure 2:**
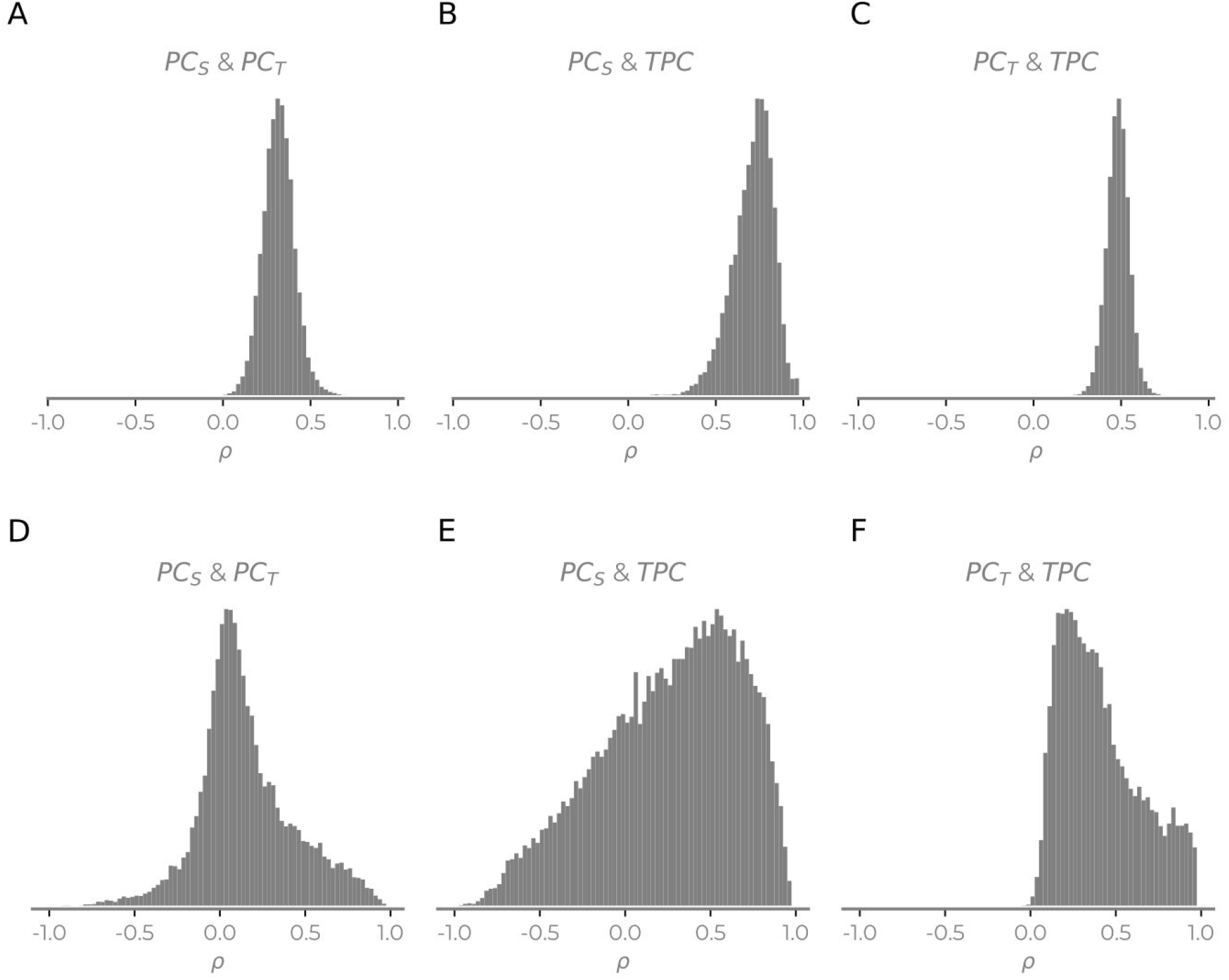
Replicating the results in Figure 5 using the MTD time-varying connectivity method and TCTC community detection method. (A-C) The histograms of the correlations of the time series for each method (histogram showing all nodes, sessions and subjects) when using the MTD method for time-varying connectivity estimates. Everything else in the main text was the same. (D-E) The histograms of the correlations of the time series for each method (histogram showing all nodes, sessions and subjects) when using the TCTC temporal community detection method for TPC and PC_T_(PC_S_ is the same as the main text). All other methodological steps are the same as the main text.

